# GenoGAM 2.0: Scalable and efficient implementation of genome-wide generalized additive models for gigabase-scale genomes

**DOI:** 10.1101/265595

**Authors:** Georg Stricker, Mathilde Galinier, Julien Gagneur

## Abstract

GenoGAM (Genome-wide generalized additive models) is a powerful statistical modeling tool for the analysis of ChIP-Seq data with flexible factorial design experiments. However large runtime and memory requirements of its current implementation prohibit its application to gigabase-scale genomes such as mammalian genomes.

Here we present GenoGAM 2.0 a scalable and efficient implementation that is 2-3 orders of magnitude faster than the previous version. This is achieved by exploiting the sparsity of the model using the SuperLU direct solver for parameter fitting, and sparse Cholesky factorization together with the sparse inverse subset algorithm for computing standard errors. Furthermore the HDF5 library is employed to store data efficiently on hard drive, reducing memory footprint while keeping I/O low. Whole-genome fits for human ChIP-seq datasets (ca. 300 million parameters) could be obtained in less than 9 hours on a standard 60-core server. GenoGAM 2.0 is implemented as an open source R package and currently available on GitHub. A Bioconductor release of the new version is in preparation.

We have vastly improved the performance of the GenoGAM framework, opening up its application to all types of organisms. Moreover, our algorithmic improvements for fitting large GAMs could be of interest to the statistical community beyond the genomics field.

## 1 Background

Chromatin immunoprecipitation followed by deep sequencing (ChIP-Seq), is the reference method for quantification of protein-DNA interactions genome-wide (1). ChIP-Seq allows studying a wide range of fundamental cellular processes such as transcription, replication and genome maintenance, which are characterized by occupancy profiles of specific proteins along the genome. In ChIP-Seq based studies, the quantities of interest are often the differential protein occupancies between experiments and controls, or between two genetic backgrounds, or between two treatments, or combinations thereof.

We have recently developed a statistical method, GenoGAM (Genome-wide Generalized Additive Model), to flexibly model ChIP-Seq factorial design experiments (2). GenoGAM models ChIP-Seq read count frequencies as products of smooth functions along chromosomes. It provides base-level and region-level significance testing. An important advantage of GenoGAM over competing methods is that smoothing parameters are objectively estimated from the data by cross-validation, eliminating ad-hoc binning and windowing. It leads to increased sensitivity in detecting differential protein occupancies over competing methods, while controlling for type I error rates.

GenoGAM is implemented as an R package based on the well-established and flexible generalized additive models (GAM) framework (3). On the one hand, it builds on top of the infrastructure provided by the *Bioconductor* software project (4). On the other hand, it uses the mgcv package (5), a general-purpose R library for fitting GAMs (6) that provides a rich functionality for GAMs with a variety of basis functions, distributions and further features for variable and smoothness selection. In its general form, the implementation for fitting a GAM minimizes a cost function using iterations whose time complexity are quadratic in the number of parameters. Moreover, the time complexity of the implementation for estimating the standard errors of the parameters, which are required for any statistical significance assessment, is cubic in the number of parameters. To allow the fitting of GAMs on complete genomes, which involves millions of parameters, we had proceeded with a tiling approach (2). Genome-wide fits were obtained by fitting models on *tiles*, defined as overlapping genomic intervals of a tractable size, and joining together tile fits at overlap midpoints. With long enough overlaps, this approximation yielded computation times linear in the number of parameters at no practical precision cost. Furthermore, it allowed for parallelization, with speed-ups being linear in the number of cores.

Nonetheless, application of the current implementation remains limited in practice to small genomes organisms such as yeast or bacteria, or to selected subsets of larger genomes. A genome-wide fit for the yeast genome (ca. 1 million parameters) took 20 hours on a 60-core server. Fits for the human genome could only be done for chromosome 22, the smallest human chromosome.

Here we introduce a new implementation of GenoGAM that is 2 - 3 orders of magnitude faster. This is achieved by exploiting the sparsity of the model and by using out-of-core data processing. The computing time for parameter and standard error estimation, as well as the memory footprint, is now linear in the number of parameters per tile. The same genome-wide fit for yeast is now obtained in 13 minutes on a standard 8-core desktop machine. Whole-genome fits for human datasets (ca. 300 million parameters each) are obtained in less than 9 hours on the same 60-core server.

Before describing the new implementation and results, we provide some necessary mathematical background.

### 1.1 GenoGAM models

In a GenoGAM model, we assume ChIP-Seq read counts *y_i_* at genomic position *x_i_* in the ChIP-Seq sample *j_i_* to follow a negative binomial distribution with mean *μ_i_* and dispersion parameter *θ*:

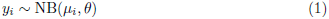

where the logarithm of the mean *μ_i_* is the sum of an offset *o_i_* and one or more smooth functions *f_k_* of the genomic position *x_i_*:

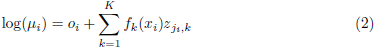

The offsets *o_i_* are predefined data-point specific constants that account for sequencing depth variations. The elements 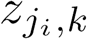 of the experimental design matrix **Z** is 1 if smooth function *f_k_* contributes to the mean counts of sample *j_i_* and 0 otherwise. A typical application is the comparison of treatment versus control samples, for which a GenoGAM model would read:

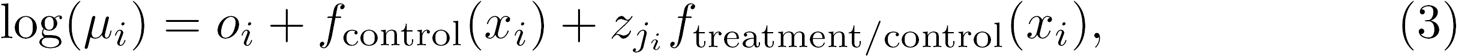

where 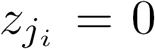 for all control sample data points and 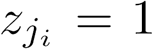 for all treatment sample data points. The quantity of interest in such a scenario is the log fold-change of treatment versus control at every genomic position *f*_treatment/control_(*x_i_*).

The smooth functions *f_k_* are piecewise polynomials consisting of a linear combination of basis functions br and the respective coefficients 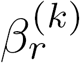:

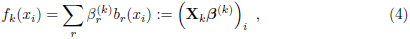

where *b_r_* are second order B-splines, which are bell-shaped cubic polynomials over a finite local support (7). The column of the *n* × *p_k_* matrix **X**_*k*_, where *p_k_* is the number of basis functions in smooth *f_k_*, represents a basis function *b_r_* evaluated at each position *x_i_*.

Typically all smooth functions have the same bases and knot positioning, implying that all **X**_*k*_ are equal to each other. Consequently, the complete design matrix **X** is the Kronecker product of the experimental design matrix **Z** and **X**_*k*_.

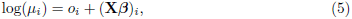

where 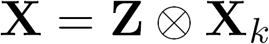 and the vector *β* is the concatenation of all *β^(k)^*.

The fitting of the parameters *β* is carried out by maximizing the negative binomial log-likelihood plus a penalty function:

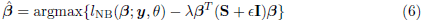

where **S** is a symmetric positive matrix that approximately penalizes the second order derivatives of the smooth functions *β*. This approach is called penalized B-splines or P-splines (8). The ϵ**I** term adds regularization on the squared values of the *β*’s, which is particularly useful for regions with many zero counts. The smoothing parameter *λ* controls the amount of regularization. Both the smoothing parameter *λ* and the dispersion parameter *θ* are considered as hyperparameters that are estimated by cross-validation (2).

Newton-Raphson methods are used to maximize Equation (6). The idea is to iteratively maximize quadratic approximations of the objective function around the current estimate. The current parameter vector *β_t_* is updated as:

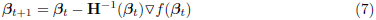

where the negative inverse Hessian **H**^-1^(*β_t_*) captures the local curvature of the objective function, and the gradient vector ▽*f*(*β_t_*) captures the local slope. The iteration stops when the change in the log-likelihood or the norm of the gradient of the log-likelihood falls below a specified convergence threshold. Because the negative binomial distribution with known dispersion parameter *θ* is part of the exponential family, the penalized log-likelihood is convex and thus convergence is guaranteed.

### 1.2 Standard error computation

For the purpose of statistical testing, variance of the smooth estimates are also needed. These are of the form:

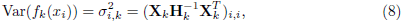

where **H***_k_* is the hessian with respect to the parameters *β*^(*k*)^ and can be simply extracted from **H**.

### 1.3 Remarks on sparsity

The Hessian **H** is computed as:

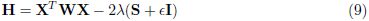

with **W** a diagonal matrix (5).

However, the number of nonzeros for each row of the design matrix **X** is at most 5 times the number of smooth functions because every genomic position *x_i_* is overlapped by 5 second order B-splines *b_r_* only. Moreover, the penalization matrix **S** only has 5 nonzeros per row, as it encodes the second-order difference penalties between coefficients of neighboring splines (8). Hence, the matrices **X** and **S**, and therefore **H**, which appears in the majority of the computations via Equations (7) and (8), are very sparse. Here we make use of the sparsity of these matrices to drastically speed up the fitting of the parameters.

## 2 Implementation

### 2.1 Workflow

Data preprocessing consists of reading raw read alignments from BAM files, centering the fragments, computing the coverage **y**, and splitting the data by genomic tiles (Figure 1). Afterwards, normalization factors for sequencing depth 6 variation are computed using DESeq2 (9). In the new version of GenoGAM we store the preprocessed data in HDF5 files (10) through the R packages HDF5Array (11) and rhdf5 (12). This allows writing in parallel as the data is being pre-processed, which reduces the memory footprint of this step. For all subsequent matrix operations the Matrix package is used, which implements routines for storage, manipulation and operations on sparse matrices (13).

**Figure 1.**
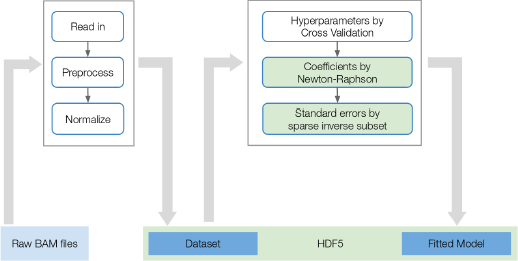
Schematic overview: Raw BAM Files are read-in, pre-processed normalized and written to hard drive in HDF5 format. Moreover, normalization factors for sequencing depth variation are computed using DESeq2 (9). The resulting object is the dataset upon which fitting is done. Then global hyper-parameters are estimated by cross-validation and for each tile coefficients are estimated via Newton-Raphson and standard errors via sparse inverse subset algorithm. The final model is written as a new object to hard drive in HDF5 format. Note, that the schematic view is a simplification: The pre-processed dataset and the fitted model are not generated in memory and written to HDF5 in the end. Instead, all HDF5 matrices are initialized on hard drive directly and the writing is done on the fly. The green marked rectangles display the areas of improvement (and the content of this paper) in this version of GenoGAM over the previous version.

Fitting GenoGAM models on tiles is achieved by the Newton-Raphson algorithm (Equation 7). This is done on few representative tiles during cross-validation in order to identify optimal hyperparameters *λ* and *θ*, and subsequently when fitting the model on the full dataset.

The variance of the smooth estimates (Equation 8) is obtained using the sparse inverse subset algorithm as detailed in a subsection below. The implementation is based on the R package sparseinv (14), which wraps relevant code from the SuiteSparse software (15). As in the previous GenoGAM model (2), fitting on different tiles is conducted in parallel. The result objects for the fits, variances and parameters are initialized prior to fitting on hard drive. This allows the processes to write results in parallel on the fly, ensuring fast computation and low memory footprint. The HDF5 storage is further optimized for reading time by adjusting HDF5 chunk size to the size of the tiles (for preprocessed count data) and chunks (for fits and variance). As HDF5 is not process-safe on R level, writing is serialized by a queuing mechanism.

The parallelization backend is provided by the R package BiocParallel. It offers an interface to a variety of backends and can be registered independently of GenoGAM. Parallelization is performed over chromosomes during the read-in process. Over tuples of folds and tiles during cross-validation process and over tiles during fitting process. Because some backends have a particular long start-up time, the use of many processes might end up dominating computation time. Specifically during cross-validation on small and limited number of regions, this might pose a problem. Therefore an optimal number of workers is automatically obtained and registered by the cross-validation function and reset on exit.

### 2.2 Newton-Raphson implementation for sparse matrices

We estimate the parameters *β* by maximizing the penalized log-likelihood using the Newton-Raphson iteration (Equation 7). Due to the sparsity of the matrices **X**, **D** and **S**, **H** is sparse and cheap to compute. The inverse is never explicitly formed. Instead the linear system is solved by a direct solver using the SuperLU library (16). Furthermore all matrices are stored in a sparse format, avoiding redundant storage of zeros.

Our new fitting algorithm differs from the one of mgcv in two ways. First, mgcv uses Iteratively Reweighted Least Squares, a Newton-Raphson method that employs the Fisher information matrix 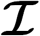, defined as the negative expectation of the Hessian **H**, instead of the Hessian in the iteration (Equation 7). However, this did not lead to any measurable differences in the fitted parameters. Second, mgcv uses QR decomposition of the design matrix **X** (5). However, general QR decomposition destroys the sparse structure of **X**. We have investigated the use of sparse QR decompositions but this was less efficient than our final implementation.

### 2.3 Variance computation using the sparse inverse subset algorithm

The hessian **H** is sparse, but its inverse, the covariance matrix **H**^-1^, usually is not. However, the variances of interest (Equation 8) can be computed using only a subset of the elements of the inverse **H**^-1^. Specifically, denoting for any matrix **A**:

- 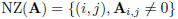 the indices of nonzero elements,
- 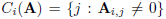 the column indices of nonzero elements for the i-th row,
- 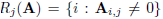 the column indices of nonzero elements for the j-th row,

then σ^2^ can be computed only using the elements 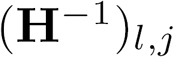, where 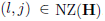. Indeed, on the one hand we have:

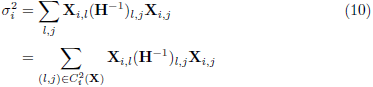

On the other hand, Equation 9 implies that 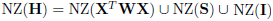. Since

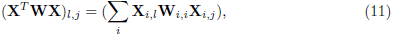

it follows that:

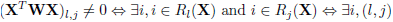

Moreover,the nonzeros of the identity matrix **I** is a subset of the nonzeros of the second-order differences penalization matrix **S** (8). Furthermore, the nonzeros of the second-order differences penalization matrix **S**, which penalizes differences between triplets of consecutive splines, is a subset of the nonzeros of **X***^T^***X**, since genomic positions overlap five consecutive splines when using second-order B-splines. Hence, 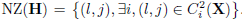. Together with Equation 10, this proves the result.

Using only the elements of **H**^-1^ that are in NZ(**H**) applies to the variance computation of any linear combinations of the *β* based on the same sparse structure of **X** or a subset of it. Hence, it applies to computing variance of the predicted value for any smooth function *f_k_*(*x*) or computing variance of the derivatives of any order *r* of any smooth 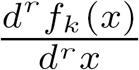.

To obtain the elements of **H**^-1^ that are in NZ(**H**), we used the sparse inverse subset algorithm (17). Given a sparse Cholesky decomposition of symmetric matrix **A** = **LL**^*T*^, the sparse inverse subset algorithm returns the values of the inverse **A**^-1^ that are nonzero in the Cholesky factor **L**. Since nonzero in the lower triangle of **A** are nonzeros in the Cholesky factor **L** (18), the sparse inverse subset algorithm provides the required elements of **H**^-1^ when applied to a sparse Cholesky decomposition of **H**. See also Rue (19) for similar ideas for Gaussian Markov Fields. To perform the sparse inverse subset algorithm, we used the R package sparseinv (14), itself a wrapper of relevant code from the SuiteSparse software (15).

Once the sparse inverse subset of the Hessian is obtained, 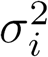 can be computed according to Equation (10) with a slight improvement: Because only the diagonal is needed from the final matrix product, the implementation does not perform two matrix multiplications. Instead, only the first product is computed, then multiplied element-wise with 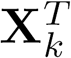 and summed over the columns.

## 3 Results

### 3.1 Leveraging the sparse data structure allows for faster parameter estimation

Figure 2 displays the comparison in fitting runtime (A) and memory usage (B) of our Newton-Raphson method versus the method underlying the previous GenoGAM version on a single core. Computation was capped at approximately 2 hours, which leads the blue line (GenoGAM 1.0) to end after around 1,100 parameters. It can be clearly seen, that exploiting the advantages of the data structures leads to improvements by 2 - 3 orders of magnitude. At the last comparable point at 1,104 parameters it took the previous method 1 hours and 37 minutes, while it was only 1 second for the Newton-Raphson method. This number increased a little bit towards the end to almost 5 seconds for 5,000 parameters.

**Figure 2.**
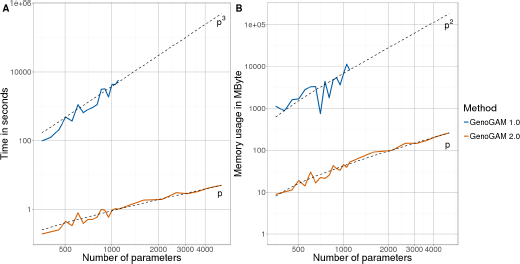
Coefficient estimation performance. (**A**) Empirical runtime for the estimation of coefficients vector *β* is plotted in log-scale against increasing number of parameters (also log-scale). The runtime is capped at around 2 hours, such that runtime of previous GenoGAM version (blue line) terminates after 1,100 parameters. The new version of GenoGAM (red line) achieves linear run-time in *p* (dotted line *p*), the number of parameters, compared to the previous cubic complexity (dotted line *p*^3^). (**B**) Memory consumption in MByte for the estimation of coefficients vector *β* is plotted against number of parameters (also log-scale). Due to the runtime cap at around 2 hours the runtime of previous GenoGAM version (blue line) does terminate after 1,100 parameters. The storage of matrices in sparse format and direct solvers avoiding full inversion keep the memory footprint low and linear in *p* (dotted line *p*) for the new GenoGAM version (red line) compared to quadratic in the previous version (blue line, dotted line *p*^2^).

Additionally, the more efficient storage of sparse matrices and the lightweight implementation reduces the overhead and memory footprint. Again at the last comparable point, the memory used by the previous method is 8 Gbyte while it is 10 52 MByte by the new method, increasing to 250 MByte at the 5,000 parameters mark. Moreover, runtime per tile drops empirically from growing cubically with the number of parameters in GenoGAM 1.0 to linearly in GenoGAM 2.0. Also, The memory footprint drops empirically from growing quadratically with the number of parameters in GenoGAM 1.0 to linearly in GenoGAM 2.0 (dashed black lines fitted to the performance data).

### 3.2 Exact σ^2^ computation by the sparse inverse subset algorithm

Alternatively to the direct computation of the inverse Hessian with consecutive computation of variance vector σ^2^, it is also possible to directly compute σ^2^. Here and hereafter the smooth function specific index *k* is dropped for simplicity. In a comment to the paper of Lee and Wand (20), a direct way to compute σ^2^ without inverting **H** was proposed by Simon Wood ((21)). The comment states, that in general, if *y* = **X**β, then

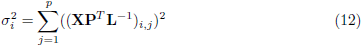

Where **P** is the permutation matrix and **L**^-1^ is the inverted lower triangular matrix resulting from Cholesky decomposition of 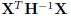.

Figure 3 shows the comparison of both methods in time and memory on a single core, with the above proposed method depicted as “indirect” (blue). While both methods have linear memory footprint, the slope of the indirect method is around four times higher. The computation time is significantly in favor of the sparse inverse algorithm. This is because for every 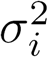 a triangular system has to be solved to obtain 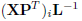. Although solving the complete system at once is faster, it had a high memory consumption when it came to increased number of parameters in our implementation. Thus the performance presented is based on batches of 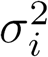 to obtain a fair trade-off between runtime and memory. Nevertheless, the difference remains around 2 orders of magnitude. Moreover runtime goes now linearly in practice for the sparse inverse subset algorithm compared to quadratically for the indirect method (dashed black lines fitted to the performance data).

**Figure 3.**
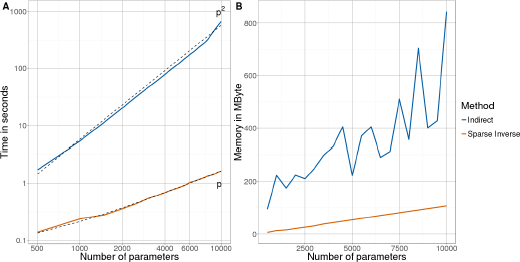
Standard error computation. (**A**) Empirical runtime for the computation of standard error vector σ^2^ is plotted in log-scale against increasing number of parameters (also log-scale). Computation based on sparse inverse subset algorithm (red line) achieves linear runtime in *p* (dotted line *p*), the number of parameters, compared to quadratic complexity (dotted line *p*^2^) of the “indirect” method (blue line). (**B**) Memory consumption in MByte for the computation of standard error vector σ^2^ is plotted against number of parameters. Though both methods achieve linear memory consumption in *p*, the slope of the “indirect” method (blue line) is around 4 times greater than of the sparse inverse subset algorithm (red line). A consequence likely due to the recursive computation of the inverse instead of solving of a triangular system.

### 3.3 Performance on human and yeast ChIP-Seq datasets

The previous version of GenoGAM could only be partially applied genome-wide for megabase-scale genomes such as the yeast genome and was impractical for gigabase-scale genomes such as the human genome. A genome-wide model fit with two conditions and two replicates each took approximately 20 hours on 60 cores (2). With computational and numerical improvement on the one side and a data model largely stored on hard drive on the other side, runtime and memory requirements have dropped significantly. Figure 4 shows the runtime performance on seven human ChIP-Seq datasets with two replicates for the IP and one or two replicates for the control. The analysis was performed with 60 cores on a cluster, the memory usage never exceeded 1.5GB per core and was mostly significantly lower. The overall results show that around 20 minutes are spend with pre-processing the data, which is largely occupied by writing the data to HDF5 files. One hour of cross-validation, to find the right hyperparameters and around 7 - 8 hours of fitting, amounting to a total runtime of 8 - 9 hours. It is also notable, that the transcription factors NRF1, MNT and FOXA1 include two controls instead of one, thus efficiently increasing the amount of data to fit by a third, but the runtime by around 40 minutes.

**Figure 4.**
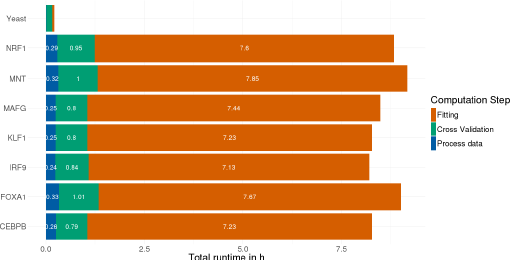
Genome-wide performance for human and yeast. The performance of GenoGAM 2.0 on seven human ChIP-Seq datasets for the transcription factors NRF1, MNT, FOXA1, MAFG, KLF1, IRF9 and CEBPB. The first three of which contain two replicates for the control, while the rest contains only one. This increases the data by around a 1/3, but the runtime by around 40 minutes, equivalent to approximately 1/11. Overall ca. 20 minutes are spent on data processing (blue), up to one hour on cross-validation (green) and 7 - 8 hours of fitting (red) amounting to a total of 8 - 9 hours runtime on 60 cores, with the snow parallel backend and HDF5 data structure. At the very top yeast runtime is shown on a regular machine with 8 cores, the multicore backend and all data kept in memory avoiding I/O to hard drive. Data processing (blue, almost not visible) takes 40 seconds, cross-validation around 9 minutes (green) and fitting 3.5 minutes (red).

Additionally, the same yeast analysis is shown by running on a laptop with 8 cores for comparison to the previous version. The total runtime is around 13 minutes with the cross-validation significantly dominating both other steps (around 9 minutes). This is due to the fact, that the number of regions used is fixed at 20, resulting in 200 model fitting runs for one 10-fold cross-validation iteration. Hence, for a small genome like the yeast genome, hyperparameter optimization may take more time than the actual model fitting. Note, that during cross-validation the only difference between human and yeast analysis is the underlying data and the parallel backend. However the runtime on yeast is only 1/6 of the runtime in human. Both factors play a role in this: First, the parallel backend in the yeast run uses the Multicore backend, allowing for shared memory on one machine. While the human run uses the Snow (simple network of workstations) backend, which needs to initiate the workers and copy the needed data first, resulting in an overall greater overhead. Second, convergence on yeast data is generally faster due to higher coverage resulting not only in less iterations by the Newton-Raphson, but also during cross-validation.

## 4 Conclusion

We have significantly improved the implementation of GenoGAM (2) on three main aspects: Data storage, coefficient estimation and standard error computation. We showed its runtime and memory footprint to scale linearly with the number of parameters per tiles. As a result, GenoGAM can be applied overnight to gigabase-scale genome datasets on a typical lab server. Runtime for mega-base genomes like the yeast genome is within minutes on a standard PC. Finally, our algorithmic improvements apply to GAMs of long longitudinal data and can therefore be relevant for a broader community beyond the field of genomics.

## 5 Availability and requirements

- Project name: GenoGAM
- Project home page: https://github.com/gstricker/fastGenoGAM
- Operating system(s): Platform independent
- Programming language: R, C++
- Other requirements: R 3.4.1 (https://cran.r-project.org/) or higher
- License: GPL-2

## 6 Declarations

### 6.1 Ethics approval and consent to participate

Not applicable

### 6.2 Consent for publication

Not applicable

### 6.3 Availability of data and materials

The datasets analyzed during the current study are available from ENCODE:

- CEBPB: https://www.encodeproject.org/experiments/ENCSR000EHE
- FOXA1: https://www.encodeproject.org/experiments/ENCSR267DFA
- IRF9: https://www.encodeproject.org/experiments/ENCSR926KTP
- KLF1: https://www.encodeproject.org/experiments/ENCSR550HCT
- MAFG: https://www.encodeproject.org/experiments/ENCSR818DQV
- MNT: https://www.encodeproject.org/experiments/ENCSR261EDU
- NRF1: https://www.encodeproject.org/experiments/ENCSR135ANT

Yeast data analyzed during this study is included in this published article from Thornton et al. (22)

### 6.4 Competing interests

The authors declare that they have no competing interests.

### 6.5 Funding

This project has received funding from the European Union’s Horizon 2020 research and innovation program under grant agreement No. 633974 (J.G. and G.S.).

### 6.6 Author’s contributions

Conceived the project and supervised the work: J.G. Developed the software and carried out the analysis: G.S., M.G. and J.G. Wrote the manuscript: G.S. and J.G.

### 6.7

#### Acknowledgements

We thank Alexander Bertram, Parham Solaimani, Martin Morgan and Hervé Pagès for support and advice during the implementation of the second version of GenoGAM, Thomas Huckle and Simon Wood for fruitful discussions and advice on sparse matrix methods.

